# The First Inhibitor of *Meso*-Diaminopimelate Biosynthesis with Antibacterial Activity Against Multi-Drug Resistant Bacteria

**DOI:** 10.1101/2022.02.10.480023

**Authors:** Tatiana P. Soares da Costa, Jessica A. Wyllie, Chamodi K. Ghardi, Mark D. Hulett, Belinda M. Abbott, J. Mark Sutton, Matthew A. Perugini

**Author notes:** Address correspondence to Tatiana P. Soares da Costa, and Matthew A. Perugini.

## Abstract

Antibiotic resistance represents one of the biggest threats to global health. While several of our current antibiotics target the peptidoglycan within the bacterial cell wall, only a fraction of its components has been explored for antibiotic development. A component that remains under-exploited is *meso*-diaminopimelate (*meso*-DAP), a constituent of the cross-linking peptide in Gram-negative bacteria. In this study, we employed a high throughput chemical screen to identify the first inhibitor of *meso*-DAP biosynthesis with antibacterial activity. Indeed, the compound was shown to have minimum inhibitory concentration values of 8–16 μg/mL against a panel of multi-drug resistant *Acinetobacter baumannii* strains, including those resistant to the last resort antibiotic carbapenem. Importantly, the compound targets the *meso*-DAP biosynthesis pathway specifically, with no off-target effects observed in human cell lines, and no resistance exhibited upon continuous treatment, under the conditions tested. Furthermore, we revealed for the first time that *meso*-DAP biosynthesis inhibition prevents biofilm formation and disrupts established biofilms in *A. baumannii*. Using a *Galleria mellonella* model, we showed that this compound improves survival rates against *A. baumannii* infection by up to 40% relative to the no treatment controls. Lastly, we determined that the inhibitor potentiates the activity of several antibiotic classes, including carbapenems. Thus, this study provides proof-of-concept that *meso*-DAP biosynthesis represents a promising target for the development of standalone antibacterial agents with a new mode of action as well as adjuvants to be used in combinatorial regimens to rejuvenate our current antibiotic arsenal to combat resistance.

**Importance:** Resistance levels to available antibiotics continues to rise, with a growing number of Gram-negative bacterial infections, in particular *A. baumannii* infections, becoming life-threatening. Despite this, there have been no new classes of antibiotics against Gram-negative bacteria introduced to the market over the last 40 years. Hence, new targets and therapeutics are urgently required to combat these clinically important pathogens. One such target is *meso*-DAP, a critical component of the cross-linking peptides in the cell walls of Gram-negative bacteria. Here, we describe the first inhibitor of bacterial *meso*-DAP biosynthesis, with antibacterial activity against multi-drug resistant Gram-negative bacterial strains, including carbapenem-resistant *A. baumannii*. We also reveal that *meso*-DAP biosynthesis inhibition affects biofilm stability and potentiates the activity of several antibiotic classes. This study highlights the need to further explore *meso*-DAP biosynthesis and other unexploited targets in the search for antibiotics with new modes of action.

## Introduction

Effective antibiotics are critical for modern medicine. However, our current options are rapidly dwindling as the prevalence of antibiotic-resistant bacteria continues to rise (1). To make matters worse, very few antibiotics with new modes of action have been introduced to the market over the last 40 years and none for Gram-negative bacteria. Therefore, it is of critical importance that we identify novel approaches to discover new classes of antibiotics to tackle resistance. The bacterial cell wall is the target of some of the most powerful antibiotics discovered to date, including carbapenems and vancomycin, two of our ‘last resort’ treatments (2, 3). These antibiotics disrupt the formation of peptidoglycan within the cell wall, resulting in cell death through osmotic lysis. The peptidoglycan is comprised of alternating disaccharide units, *N*-acetylglucosamine and *N*-acetylmuramic disaccharide, cross-linked by short peptide chains (4). Despite the clinical success of inhibiting peptidoglycan biosynthesis, only a fraction of its components has been explored as targets for antibiotic development. A component that remains under-exploited is *meso*-diaminopimelate (*meso*-DAP), a constituent of the cross-linking peptide in Gram-negative bacteria (4).

The biosynthesis of *meso*-DAP occurs via the DAP pathway that is exclusively found in bacteria and plants (5, 6) (Figure 1). The pathway begins with the condensation of pyruvate and aspartate semialdehyde (ASA) to form 4-hydroxy-2,3,4,5-tetrahydrodipicolinic acid (HTPA), which is catalysed by 4-hydroxy-tetrahydrodipicolinate synthase, commonly known as dihydrodipicolinate synthase (DHDPS, E.C. 4.3.3.7) (7) (Figure 1). The product is reduced to 2,3,4,5-tetrahydrodipicolinate (THDP) by dihydrodipicolinate reductase (DHDPR, E.C. 1.17.1.8) (8) (Figure 1). The pathway then diverges into four sub-pathways. In Gram-negative bacteria, L,L-DAP is converted to *meso*-DAP by diaminopimelate epimerase (DAPEP, EC 5.1.1.7) via the succinyl sub-pathway (9) (Figure 1). In addition to being used in the assembly of the peptidoglycan in Gram-negative bacteria, *meso*-DAP can also be irreversibly decarboxylated by diaminopimelate decarboxylase (DAPDC, E.C. 4.1.1.20) to form the amino acid, L-lysine (lysine). Lysine regulates flux through the pathway by binding allosterically to DHDPS and inhibiting the enzyme (10, 11) (Figure 1). Thus, DHDPS catalyses the rate-limiting step of the DAP pathway (12, 13).

**Figure 1:**
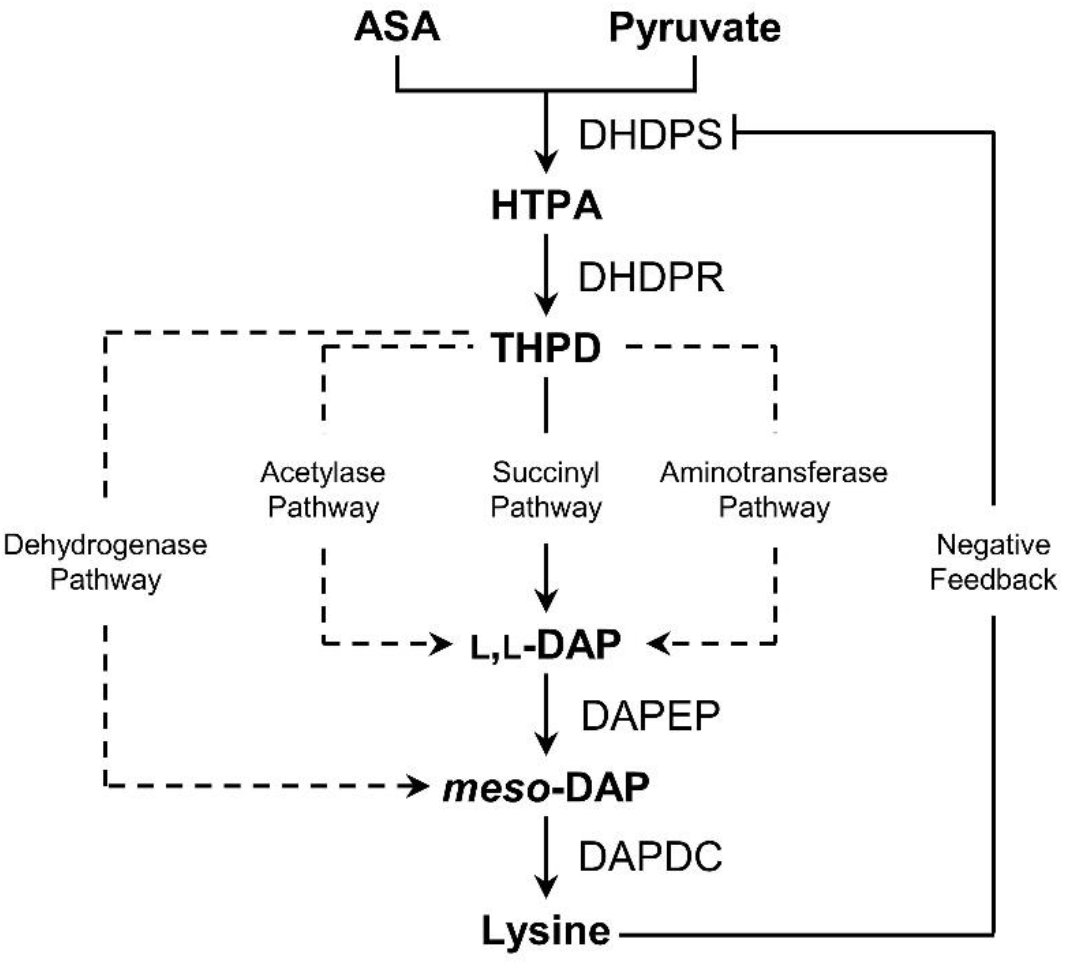
Diaminopimelate pathway in bacteria. The pathway begins with the condensation reaction between L-aspartate semialdehyde (ASA) and pyruvate to form 4-hydroxy-2,3,4,5-tetrahydrodipicolinic acid (HTPA) catalysed by dihydrodipicolinate synthase (DHDPS). Dihydrodipicolinate reductase (DHDPR) then catalyses the reduction of HTPA to produce 2,3,4,5-tetrahydrodipicolinte (THDP). In Gram-negative bacteria, THDP is subsequently converted via the succinyl sub-pathway to L,L-DAP, which is converted to *meso*-DAP by diaminopimelate epimerase (DAPEP). Finally, *meso*-DAP can be irreversibly decarboxylated by diaminopimelate decarboxylase (DAPDC) to L-lysine (lysine), which allosterically regulates the pathway via inhibition of DHDPS.

Due to the central role of DHDPS in cell wall biosynthesis in Gram-negative bacteria as well as bacterial protein biosynthesis, it is not surprising that its encoding gene has been shown to be essential (14–17). Given its essentiality to pathogenic bacteria and absence in humans, DHDPS is considered a promising antibiotic target (16). Indeed, a number of bacterial DHDPS inhibitors have been reported that mimic the enzyme’s substrates or product, but none have been shown to display antibacterial activity (18–24). Other enzymes in the DAP pathway have also been explored for antibiotic development but no inhibitor with antibacterial activity has been published to date (25, 26). The focus of this study was to identify the first antibacterial compound that targets *meso*-DAP biosynthesis. Specifically, we set out to identify inhibitors of DHDPS from the high priority pathogen, *Acinetobacter baumannii*.

*A. baumannii* is an opportunistic Gram-negative pathogen responsible for severe infections in the blood, urinary tract, lungs and surgical sites (27). Alarmingly, numerous multi-drug resistant (MDR) *A. baumannii* isolates have been reported, particularly in hospitals, including those resistant to carbapenem (28). The continual increase in the rates of emergence and dissemination of MDR *A. baumannii* strains is of great public health concern and the World Health Organization (WHO) has listed carbapenem-resistant *A. baumannii* in the highest priority group for which there is a critical need for the development of new antibiotics (28).

The structure for *A. baumannii* DHDPS (AbDHDPS) has been solved (PDB: 3PUD). It exists as a dimer, structurally reminiscent of the tight-dimer unit of DHDPS seen in other organisms (Figure 2A) (29). The active site is located within the (β/α)_8_-barrel, which is highly conserved across DHDPS enzymes, and the allosteric cleft is found in the dimer interface (Figure 2C) (8, 30–33). We have recently characterised the kinetic parameters of the recombinant AbDHDPS enzyme, including affinities for the substrates, pyruvate and ASA, to allow for screening of inhibitors *in vitro* (34). The structural and functional characterisation of this enzyme offers an excellent platform for inhibitor identification in the pursuit of antibacterial agents against *A. baumannii*.

**Figure 2:**
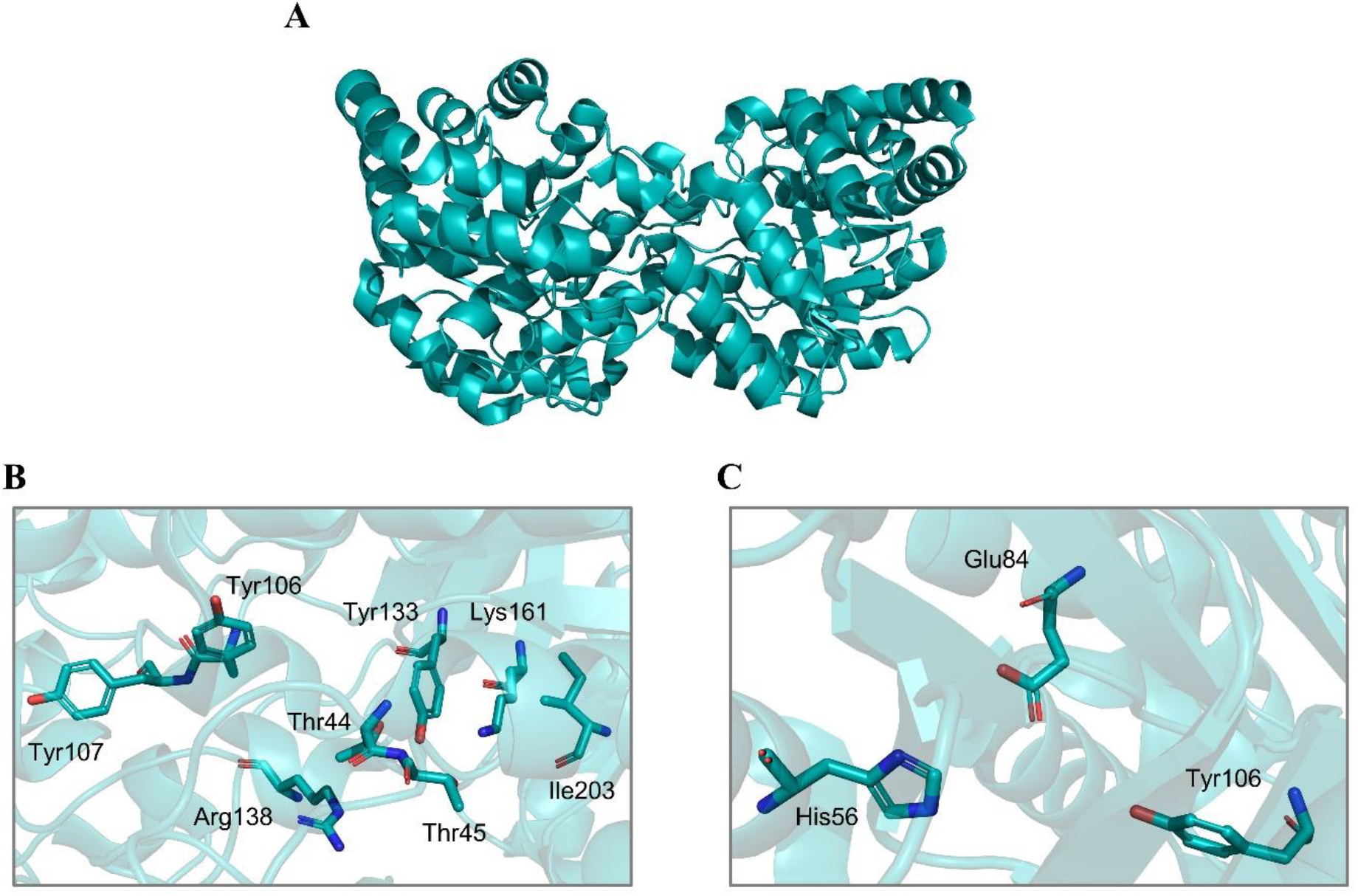
Structure of *A. baumannii* DHDPS enzyme. (A) Cartoon structure of *A. baumannii* (Ab) DHDPS (PDB: 3PUD) in the unliganded form illustrating the dimer conformation. (B) Cartoon structure of AbDHDPS, with residues within the active site shown as sticks. (C) Cartoon structure of AbDHDPS, with residues important for lysine binding and allosteric regulation shown as sticks. Residues are coloured by nitrogen (blue) and oxygen (red). Images were generated using PyMOL v 2.2 (Schrödinger).

In this study, we employed a high throughput chemical screen to identify the first inhibitor of the DAP pathway with antibacterial activity, *N*’-phenylmethanesulfonohydrazide (PMSH), through the inhibition of AbDHDPS. Specifically, this compound has low micromolar potency against the recombinant enzyme and displays antibacterial activity against a panel of MDR *A. baumannii* strains, including those resistant to carbapenem. Antibacterial activity is also observed against other high priority Gram-negative bacteria, corroborating our proposal that PMSH binds in the highly conserved DHDPS active site. Importantly, we confirmed that PMSH targets the DAP pathway specifically in bacteria by using supplementation assays and that no effect on the viability of human cell lines is observed. Furthermore, no resistance was observed upon continuous exposure of PMSH, at least under the conditions tested, and we showed for the first time that *meso*-DAP biosynthesis inhibition prevents biofilm formation and disrupts established biofilms in *A. baumannii*. Using a *Galleria mellonella* model, we demonstrated that this compound improves survival rates against *A. baumannii* infection by up to 40% relative to a vehicle control. Lastly, we determined that PMSH potentiates the activity of several antibiotic classes, including carbapenems.

## Results

### Identification of DHDPS inhibitor

A high throughput screen of a library of 87,648 compounds was conducted against recombinant DHDPS enzyme by the Walter and Eliza Hall Institute High Throughput Chemical Screening Facility (Melbourne, Australia) as described previously (35). The *o*-aminobenzaldehyde (*o*-ABA) colourimetric assay was used to assess DHDPS activity via the formation of a purple chromophore that can be measured at 520–540 nm (36). The most promising compound from the screen was *N*’-phenylmethanesulfonohydrazide (PMSH) (Figure 3A). Subsequently, the inhibitory activity of PMSH against recombinant AbDHDPS was quantitated using a DHDPS-DHDPR coupled assay (37). Different concentrations of PMSH were titrated to the assay with substrates fixed at previously determined *K*_M_ values (34), resulting in an *IC*_50_ value of 35 µM (Figure 3B).

**Figure 3:**
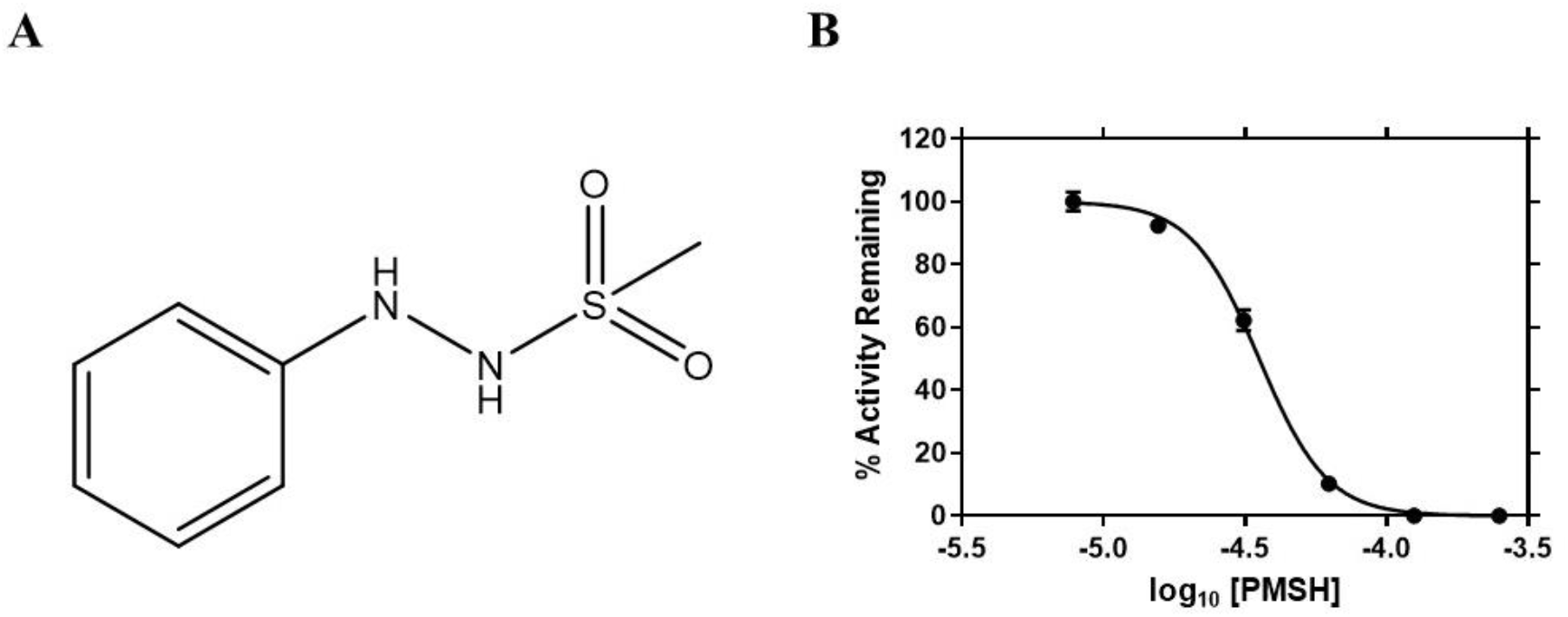
DHDPS inhibitor, PMSH. (A) Chemical structure of the compound, PMSH. (B) Dose responses of PMSH against recombinant AbDHDPS. Initial enzyme rates were normalised against a vehicle control (1% (v/v) DMSO). Normalised data (% activity remaining) is plotted as a function of log_10_[inhibitor] and fitted to a nonlinear regression model (solid line) (R^2^ = 0.99). Data represents mean ± S.D. (n = 3).

### Mode of binding of PMSH

Next, we set out to assess the mode of binding of PMSH. Attempts at crystallising the DHDPS-PMSH complex were unsuccessful. Therefore, *in silico* docking studies were performed to identify the potential residues involved in the binding of PMSH to AbDHDPS. This revealed that PMSH binds within the highly conserved active site of AbDHDPS, involving polar interactions with Tyr133, Arg138 and Asn248 (Figure 4).

**Figure 4:**
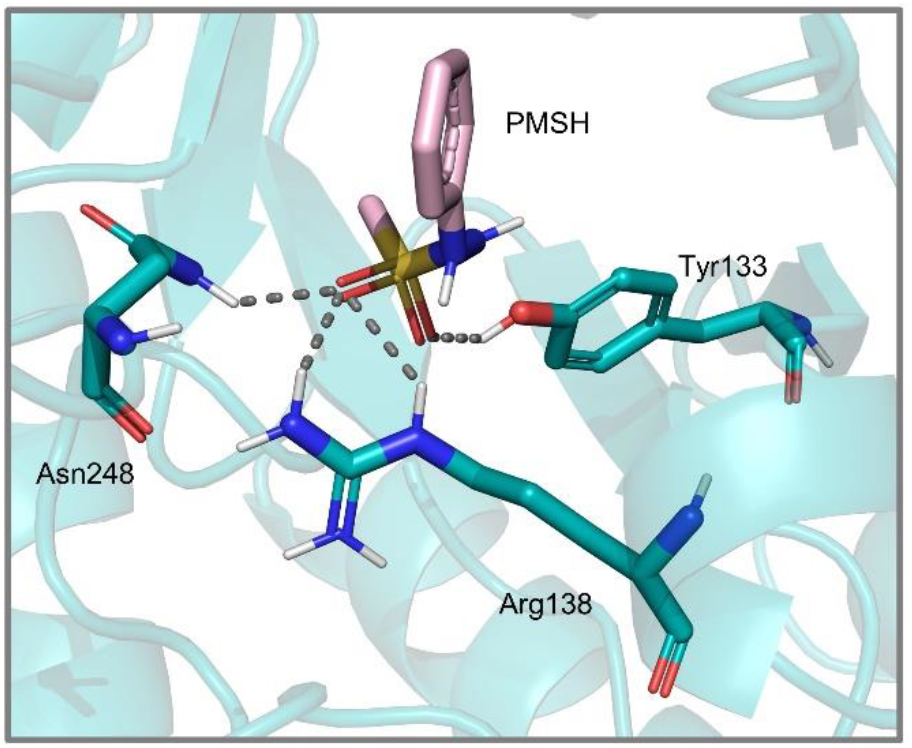
Molecular docking of PMSH to AbDHDPS. Proposed binding interactions (grey, dashed) of PMSH (pink, stick) to the active site of AbDHDPS (PDB: 3PUD) (teal, cartoon). Residues involved in binding are shown as sticks and coloured by nitrogen (blue), oxygen (red) and sulfur (yellow). Image was generated using PyMOL v 2.2 (Schrödinger)

### Antibacterial activity and specificity

The antibacterial activity of PMSH was assessed by measuring the minimum inhibitory concentration (MIC), which is defined as the lowest concentration required to prevent visible bacterial growth. These studies revealed that PMSH resulted in MIC values in the range of 8–16 μg/mL against clinical panels of drug sensitive and MDR *A. baumannii*, including strains resistant to the carbapenem, meropenem (Table 1). The lack of significant difference in susceptibility between drug sensitive and MDR strains implies that our compound is not affected by pre-existing mechanisms that mediate resistance to some of the current antibiotics used to treat *A. baumannii*. Given that PMSH is predicted to bind in the highly conserved DHDPS active site, its activity was also tested against other Gram-negative bacteria, resulting in MICs of 64 μg/mL for representative *Pseudomonas aeruginosa* isolates (PAO1 and NCTC 13437) and 64–128 μg/mL for *Escherichia coli* isolates.

**Table 1:** Minimum inhibitory concentration (MIC) values for PMSH against drug sensitive and multi-drug resistant *A. baumannii* (*A. baylyi* ADP1) strains.

| *Acinetobacter* strain | Drug resistance profile | MIC (µg/mL) |
| --- | --- | --- |
| ATCC17978 | - | 16 |
| AYE | GEN, CIP, TZP, LVX, AMK | 16 |
| W1 | - | 16 |
| ADP1 | GEN, CIP, TZP, IMP, LVX, AMK, MEM | 8 |
| NCTC13424 | GEN, CIP, TZP, LVX, AMK, MEM | 16 |
| UKA7 | GEN, CIP, LVX, AMK | 16 |
| UKA2 | GEN, CIP, TZP, IMP, LVX, AMK, MEM | 16 |
| NCTC13302 | GEN, CIP, TZP, LVX, MEM | 16 |

GEN = gentamycin, CIP = ciprofloxacin, TZP = tazobactam, LVX = levofloxacin, AMK = amoxicillin, MEM = meropenem.

Focussing on *A. baumannii*, the mechanism of antibacterial action was assessed with supplementation assays, where *meso*-DAP was added to *A. baumannii* ATCC17978 cultures in the presence and absence of PMSH at 16 μg/mL (1 × MIC). Addition of *meso*-DAP completely alleviated antibacterial activity, strongly suggesting that the target is indeed within the DAP pathway (Figure 5A). Data is expressed as percentage viability in relation to a culture of the same strain treated with vehicle. We also assessed the propensity for ATCC17978 and AYE to develop resistance to PMSH. Specifically, bacterial cultures were grown in increasing concentrations of PMSH from 0.25 × MIC up to 4 × MIC. Interestingly, no visible growth was observed when PMSH was added at 2 × MIC and greater, with no colonies observed in tryptic soy agar plates for cultures treated with 4 × MIC. Therefore, no resistant *A. baumannii* strains were observed upon treatment with PMSH under the conditions used in this study.

**Figure 5:**
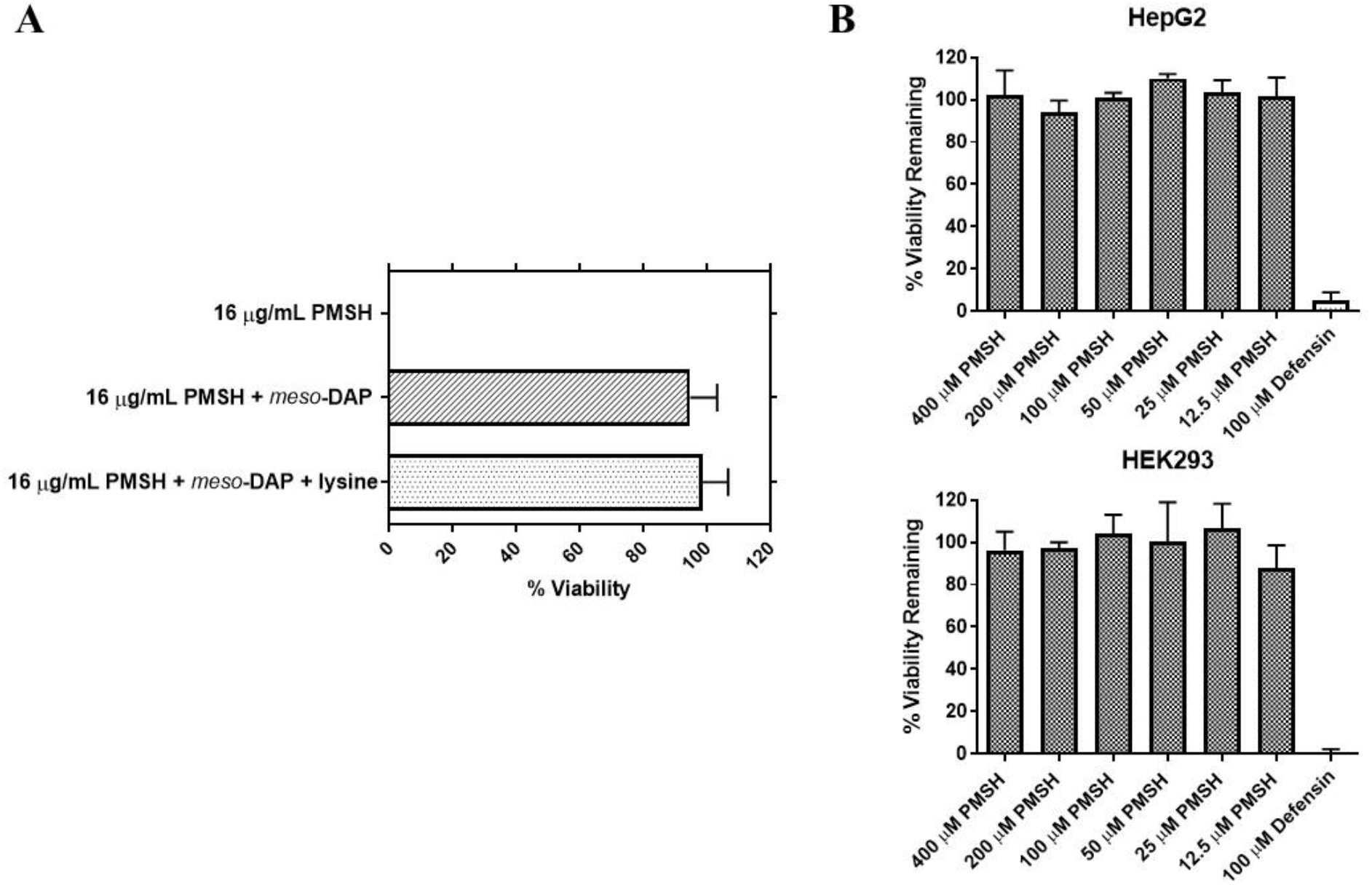
Specificity of PMSH. (A) Assessment of the mode of action of PMSH in *A. baumannii* ATCC17978 by performing supplementation experiments in the absence and presence of 500 µM *meso*-DAP with/without 500 µM lysine. Data represents mean ± S.D. (n = 3). (B) Toxicity of PMSH compared to the positive control defensin, assessed against HepG2 and HEK293 human cell lines using the MTT assay. Data were normalised against a vehicle control (1% (v/v) DMSO) and plotted against inhibitor concentration. Data represents mean ± S.D. (n = 3).

The toxicity of PMSH was addressed in a cell culture model using the human cell lines, HepG2 and HEK293. These studies showed that there was no significant reduction in viability when the cell lines were treated with PMSH even up to 400 µM (Figure 5B).

### Biofilm prevention and eradication

*A. baumannii* is notorious for developing resistance to many antibiotics due to the formation of biofilms (38). Given the involvement of *meso*-DAP in cell wall formation, this study assessed for the first time the effect of its depletion on the ability of *A. baumannii* to form biofilms, as well as its impact on established biofilms. Crystal violet staining was used to quantify *A. baumannii* biofilm mass remaining upon treatment with PMSH at varying concentrations relative to the no treatment control (DMSO) (Figure 6). Even at concentrations as low as 1 × MIC, i.e., 16 μg/mL, PMSH was able to prevent biofilm formation completely in ATCC17978 and AYE (Figure 6A). Indeed, the resulting minimum biofilm prevention concentration (MBPC) values were 16 μg/mL for both strains, the same value as obtained for the MIC measuring planktonic growth. When examining PMSH’s ability to eradicate established biofilm, we observed a concentration dependent response with both *A. baumannii* strains (Figure 6B). The effect is more pronounced on strain ATCC17978, which is a relatively poor biofilm former (38), resulting in a minimum biofilm eradication concentration (MBEC) of 64 μg/mL, relative to a MBEC of 256 μg/mL for AYE, which forms better biofilms.

**Figure 6:**
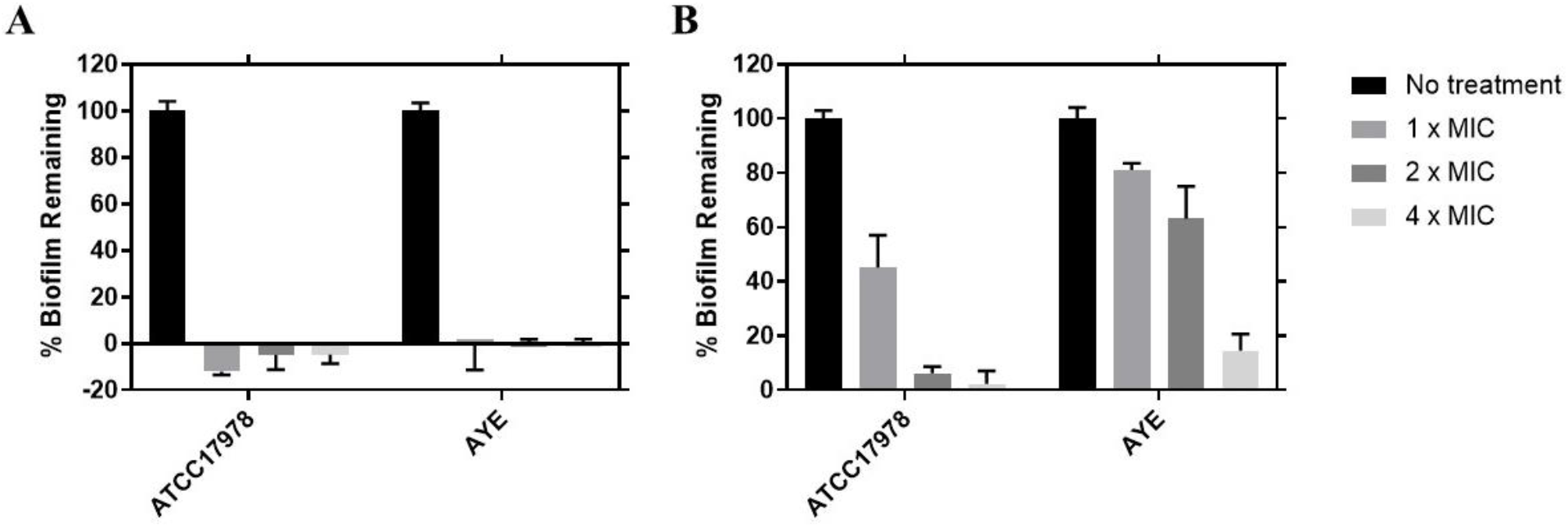
Effect of PMSH on biofilm formation and disruption in *A. baumannii* ATCC17978 and AYE. (A) Ability of PMSH to prevent biofilm formation at 1 × MIC, 2 × MIC and 4 × MIC values (grey) relative to the no treatment control (black). Data represents mean ± S.D. (n = 3). (B) Ability of PMSH to disrupt established biofilm at 1 × MIC, 2 × MIC and 4 × MIC values (grey) relative to the no treatment control (black). Data represents mean ± S.D. (n = 3).

### *In vivo* efficacy in larvae

Given the promising *in vitro* activity and specificity of PMSH, we set out to assess its efficacy *in vivo* using *G. mellonella*, which is a well-established model to study bacterial virulence and for assessing antibiotic efficacy (38, 39). *G. mellonella* larvae were infected with two clinical strains of *A. baumannii* to evaluate the *in vivo* efficacy of PMSH at a standard clinical antibiotic dose of 20 mg/kg. This was assessed by measuring larvae survival rates over a 120-hr period post injection (Figure 7). The strains used were ATCC17978, a drug sensitive *A. baumannii* strain, and AYE, which is resistant to gentamycin, ciprofloxacin, tazobactam, levofloxacin and amoxicillin (Table 1). After a 24-hr inoculation with ATCC17978, 0% survival was observed with the no treatment control, whereas addition of PMSH resulted in a survival rate of 40%, which was maintained over the 120-hr period (Figure 7A). Upon inoculation of AYE with no treatment, we observed a gradual decrease in larvae survival rate, with a 30% survival rate measured at the 120-hr time point. Addition of PMSH improved the survival rate by 20–30% relative to the no treatment control across all time intervals, resulting in a >60% survival rate at the 120-hr time point (Figure 7B).

**Figure 7:**
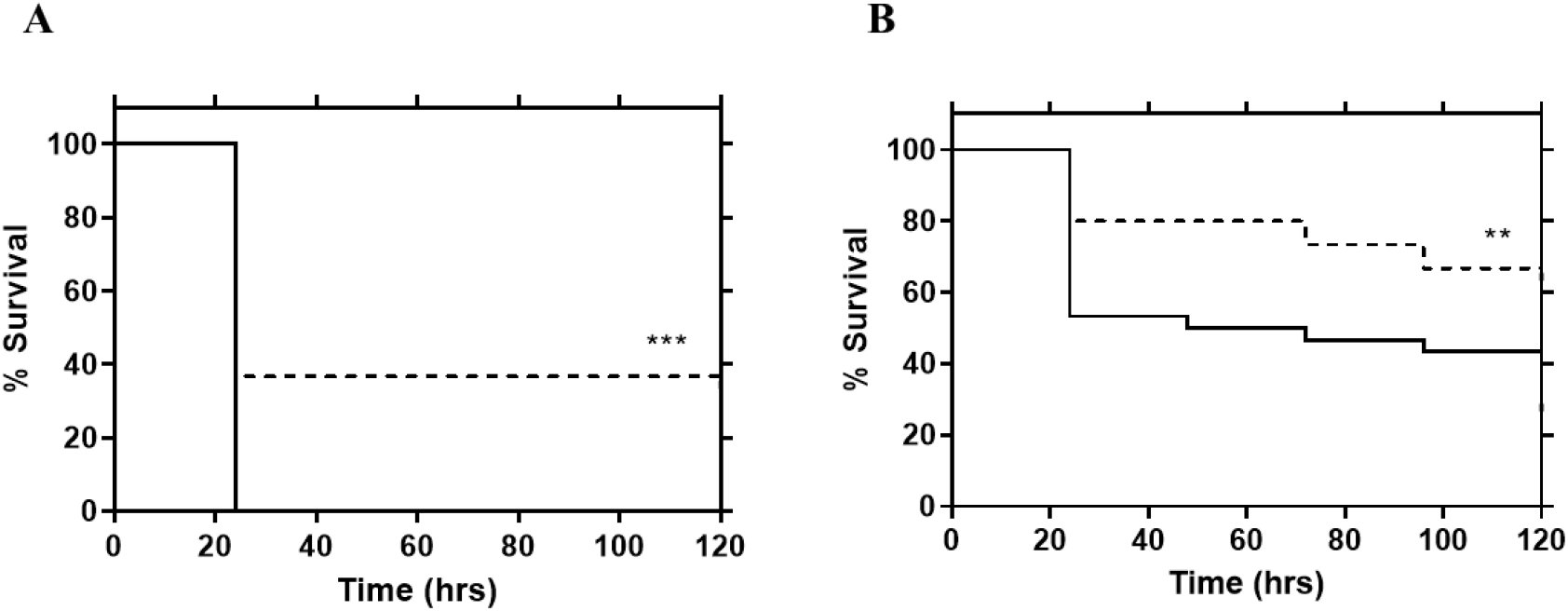
*Galleria mellonella* virulence assay. Assessment of survival of *A. baumannii* strains ATCC17978 (A) and AYE (B) before (solid line) and after (dashed line) treatment with 20 mg/kg PMSH in *G. mellonella* larvae (N = 30) up to 120 hours. The observed effects were statistically significant based on the Mantel–Cox test. ***=P≤0.001, **=P≤0.01.

### Synergistic response with current antibiotics

Lastly, we assessed the synergistic response against *A. baumannii* strains ATCC17978 and AYE with PMSH in combination with different antibiotic classes, including ceftazidime (cephalosporin), ciprofloxacin (fluoroquinolone), meropenem (carbapenem), rifampicin (rifampin) and tobramycin (aminoglycoside). The calculated fractional inhibitory concentration index (FICI) values from the checkerboard synergy assays are shown in Table 2. Interestingly, PMSH was shown to synergise with all antibiotic classes tested, yielding FICI values between 0.1741–0.3994 against both drug sensitive and drug resistant strains (Table 2). The tobramycin–PMSH combination against ATCC17978 had the lowest FICI values (Table 2). The potentiation of activity of multiple antibiotic classes is consistent with the expected mode of action of PMSH, which is most likely through the disruption of peptidoglycan synthesis, resulting in increased membrane permeability.

**Table 2:** Synergistic responses between DHDPS inhibitor and antibiotics. The calculated fractional inhibitory concentration index (FICI) was used to assess nature of interactions in combinatorial treatments. FICI < 0.5 = synergism.

| Combinatorial treatment | FICI |  |
| --- | --- | --- |
|  | ATCC17978 | AYE |
| PMSH + ceftazidime | 0.3959 | 0.3016 |
| PMSH + ciprofloxacin | 0.3478 | 0.3033 |
| PMSH + meropenem | 0.3994 | 0.3738 |
| PMSH + rifampicin | 0.3646 | 0.3158 |
| PMSH + tobramycin | 0.1741 | 0.3025 |

## Discussion

Despite the increasing prevalence of antibiotic-resistant bacteria, there have been few antibiotics with novel modes of action entering the market over the last 40 years (1). Peptidoglycan biosynthesis is a rich source of antibiotic targets, however only a fraction of its components has been explored commercially. One such under-exploited component is *meso*-DAP biosynthesis, which occurs via the DAP pathway, with the enzyme DHDPS catalysing the first committed step (4). Due to its essentiality to bacteria and absence in humans, DHDPS is considered a promising antibiotic target (16). However, whilst a number of bacterial DHDPS inhibitors have been reported to date, none have demonstrated antibacterial activity (19–24).

Here, we describe the discovery and characterisation of the first inhibitor of the DAP pathway with antibacterial activity, PMSH, via inhibition of DHDPS. When screened against clinical panels of drug sensitive and MDR *A. baumannii*, PMSH has comparable activity across all strains indicating that the compound is not affected by pre-existing mechanisms that mediate resistance to current antibiotics. Activity was also observed against other Gram-negative bacteria. The broad-spectrum activity of PMSH corroborates with the molecular docking studies, which suggested the inhibitor binds within the highly conserved DHDPS active site. PMSH was shown to be specific for the DAP pathway, reducing the propensity for potential off-target effects. Indeed, there was no observed effect of the inhibitor on the viability of human cell lines even at high micromolar concentrations. Continuous exposure of *A. baumannii* to increasing inhibitor concentrations failed to generate resistance. Additionally, AbDHDPS inhibition prevents biofilm formation and disrupts established biofilms, demonstrating for the first time that *meso*-DAP biosynthesis is critical for biofilm stability. Furthermore, in a *G. mellonella* model, PMSH significantly improves larval survival against both drug sensitive and MDR *A. baumannii* infections. Lastly, we determined that PMSH potentiates the activity of several antibiotic classes, including carbapenems, consistent with the expected mode of action of peptidoglycan biosynthesis disruption, which would result in increased membrane permeability.

It may be possible to build on the structure of PMSH to involve tighter and more extensive binding within the active site of DHDPS, improving antibacterial activity against Gram-negative bacterial pathogens. Moreover, DHDPS is one of many enzymes involved in *meso*-DAP biosynthesis, with the remaining enzymes yet to be exploited as potential antibacterial targets. Given the ability of PMSH to potentiate the activity of other classes of antibiotics, it stands to reason that a dual targeted approach to *meso*-DAP biosynthesis inhibition could be possible. Thus, this study provides proof-of-concept that DHDPS inhibitors could represent an emerging novel class of antibacterials and highlights the importance of exploring DAP pathway enzymes as potential antibiotic targets for the treatment of infections caused by high priority Gram-negative bacterial pathogens.

## Materials & Methods

### Production of recombinant *A. baumannii* DHDPS enzyme

The synthetic gene coding for the *A. baumannii* protein with a 6 × His-tag were codon optimised, synthesised and ligated into a pET28a expression vector by Bioneer Pacific (Bioneer Pacific, Kew East, Victoria, Australia). Recombinant protein was expressed in *Escherichia coli* BL21 (DE3) cells and purified using immobilised metal affinity chromatography (IMAC) as previously described (40–42). Briefly, recombinant protein was expressed upon addition of 0.5 mM isopropyl β-D-1-thiogalactopyranoside in Luria-Bertani (LB) broth at 16 °C for 18 hrs. Cells were harvested via centrifugation at 5000 × g at 4 °C and sonicated in 20 mM Tris, 150 mM NaCl, 20 mM imidazole, pH 8.0 using a Vibra Cell VC40 (Sonics and Material, Newtown, Connecticut, USA). Recombinant His-tagged proteins were purified using a 5 mL IMAC column (Bio-Rad Laboratories, Gladesville, New South Wales, Australia) and stored in 20 mM Tris, 150 mM NaCl, pH 8.0.

### High throughput chemical screen

A high throughput screen of a library of 87,648 compounds was conducted against recombinant DHDPS enzyme by the Walter and Eliza Hall Institute High Throughput Chemical Screening Facility (Melbourne, Australia) as described previously (35). Briefly, the *o*-ABA colourimetric assay was employed to assesses DHDPS activity via the formation of a purple chromophore that can be measured at 520-540 nm (36).

### Synthesis of *N*’-phenylmethanesulfonohydrazide (PMSH)

To a mix of freshly distilled aniline (1.0 mmol, 94 µL) and 1 M hydrochloric acid (5 mL) at 0 °C was added a cold 1 M aqueous solution of sodium nitrite (1.0 mmol, 1 mL) dropwise, maintaining the temperature at 0 °C, and the resultant mixture stirred for 5 mins. Addition of a cold solution of sodium metabisulphite (2.4 mmol, 456 mg) in water (3 mL) was then undertaken while maintaining the temperature at 0 °C and the reaction was stirred for a further 5 mins. The reaction was heated overnight with an oil bath at 100 °C. After cooling to room temperature, methanesulfonyl chloride (1.0 mmol, 77.4 µL) was added and the solution was neutralised by the addition of sodium carbonate. Stirring overnight at room temperature afforded the desired compound as a white precipitate (38 mg, 43%). δH (400 MHz, CDCl_3_) 8.93 (s, 1H, NH), 7.90 (s, 1H, NH), 7.14 (t, J = 7.6 Hz, 2H, ArH), 6.85 (d, J = 7.6 Hz, 2H, ArH), 6.71 (t, J = 7.6 Hz, 1H, ArH), 2.94 (s, 3H, CH_3_). δC (400 MHz, CDCl_3_) 147.1, 129.2, 120.9, 113.4, 38.8. LRMS (ESI): m/z 208.9 (M+Na)^+^, 187.0 (M+H)^+^.

### Enzyme coupled assay

The DHDPS-DHDPR coupled assay was used to quantify DHDPS activity as previously described (40, 43) by measuring the oxidation of NADPH at 340 nm. Assays were conducted in a Cary 4000 UV/Vis spectrophotometer (Varian, Mulgrave, Victoria, Australia) with substrates fixed at the previously determined *K*_M_ values (34). AbDHDPS enzyme and reactions were incubated at 37 °C for 12 mins before initiation with ASA. Initial velocity data were normalised against a vehicle (DMSO) control and analysed using Equation 1 (log(inhibitor) vs. normalized response - variable slope) using Prism Software Version 8 (Graphpad, San Diego, CA, USA). Dose responses were performed with three technical replicates for each concentration of compound.

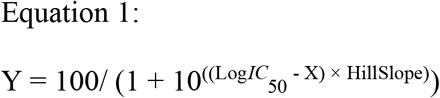

Where Y is the normalised rate, log*IC*_50_ is the logarithmic concentration of ligand resulting in 50% activity, X is the concentration of ligand, and Hill Slope is the steepness of the curve.

### *In silico* docking

The mode of binding of PMSH was assessed by performing molecular docking simulations using the AutoDock Vina tool (44) compiled in PyRx (45). The three-dimensional structure of AbDHDPS (PDB: 3PUD) was used as the macromolecule. The search space encompassed the whole of the modelled structure with the following dimensions in Å: centre (x, y, z) = (−40.28, -22.97, -20.79), dimensions (x, y, z) = (78.05, 57.42, 54.01). The docking simulations were run at an exhaustiveness of 8. The proposed models were ranked based on calculated binding affinities.

### Antibacterial assays

The MIC for PMSH was determined against a panel of *A. baumannii* strains using a broth microdilution method as described previously (46–48). Specifically, an inoculum of 1 × 10^5^ colony forming units per mL was used and the testing conducted using tryptic soy broth in 96-well plates. Growth was assessed after incubation at 37 °C for 20 hrs by measuring the absorbance at 600 nm. For supplementation assays, antibacterial assays were performed for PMSH against a panel of *A. baumannii* strains as described above, but with the addition of 500 µM *meso*-DAP or 500 µM *meso*-DAP + 500 µM L-lysine to all wells. The ability of growth recovery was also assessed relative to the no supplementation controls. All antibacterial assays were performed in 3 biological replicates.

### Toxicity assays

The viability of the human cell lines, HepG2 and HEK293, was assessed using the MTT viability assay as previously described (49–51). Briefly, the cells were suspended in Dulbecco-modified Eagle’s medium containing 10% (v/v) fetal bovine serum and then seeded in 96-well tissue culture plates at 5,000 cells per well. After 24 hrs, cells were treated with 50 – 400 µM of PMSH, such that the DMSO concentration was consistent at 1% (v/v) in all wells. As a positive control, cells were treated with the cytotoxic defensin protein at 100 µM (52). After treatment for 48 hrs, MTT cell proliferation reagent was added to each well and incubated for 3 hrs at 37 °C. The percentage viability remaining reported is relative to the vehicle control of 1% (v/v) DMSO. Assays were performed in triplicate.

### Resistance studies

ATCC17978 and AYE strains were continuously exposed to PMSH using the extended gradient MIC method from 0.25 × MIC up to 4 × MIC (53). Experiments were carried out in triplicate, with a 1% (v/v) DMSO control performed concurrently. Growth was assessed visually and by plating cultures on tryptic soy agar plates.

### Biofilm prevention and disruption assays

Cultures of ATCC17978 and AYE were grown overnight at 37 °C. To assess biofilm prevention, dilutions of PMSH were set up in Nunc-Immuno MicroWell plates to give final concentrations of 1 × MIC, 2 × MIC and 4 × MIC for each bacterial strain. Bacteria were added to give a final OD_600_ of 0.01, peg lids added, and the plates incubated for 24 hrs at 37 °C. The medium and planktonic cells were removed, and the biofilms were washed twice with 1 × PBS using gentle pipetting. Biofilms were fixed for 60 mins at 80 °C, and stained using 0.2% (v/v) crystal violet solution for 15 mins at room temperature as previously described (39, 54). The plates were washed, air-dried and de-stained with absolute ethanol for 20 mins at room temperature. The absorbance at 570 nm was measured using a SpectraMax® M5e Multi-Mode Microplate Reader. For the biofilm disruption assay, bacteria were added to Nunc-Immuno MicroWell plates and peg lids at a final OD_600_ of 0.01, and incubated for 24 hrs at 37 °C. Subsequently, PMSH at 1 × MIC, 2 × MIC and 4 × MIC were added to plates and incubated for a further 24 hrs. Biofilm biomass was assessed using crystal violet as described for the biofilm prevention assay. Experiments were conducted with three technical replicates using 1% (v/v) DMSO as the no treatment control for comparison to allow us to calculate MBFC and MBEC values.

### *G. mellonella* infection model

The *in vivo G. mellonella* survival assay was conducted as previously described (38, 55). The larvae used were about 2-3 cm long and weighed around 300 mg. Bacteria from an overnight culture of *A. baumannii* strains ATCC17978 or AYE were diluted to an OD_600_ of 0.03 in 1 × PBS and a microfluidic syringe pump was used to inject 10 μL of suspension into *G. mellonella* larvae. Injections were performed into the haemolymph of 10 larvae per treatment via the foremost left proleg. Control larvae were injected with 10 μL of 1 × PBS in order to measure any potential lethal effects of the injection procedure as well as to measure the tolerance of the larvae to the treatments. After a 30 min recovery, larvae were injected with PMSH at 20 mg/kg or 5% (v/v) DMSO. Larvae were then incubated statically at 37 °C and observed daily for 7 days post-injection. Each treatment group had 10 larvae and experiments were performed in three biological replicates. Data were analysed using the Mantel–Cox method using Prism Software Version 8 (GraphPad, San Diego, CA, USA).

### Antibacterial combinatorial treatments

Combinatorial treatments of PMSH with antibiotics were carried out using a checkerboard method (49, 56). The MIC values of compound combinations were determined after an overnight incubation at 37 °C. The nature of the interaction between PMSH and antibiotics was assessed based on the calculated FICI. The FICI of each combination was calculated as a ratio of the MIC of compound A (or B) when used in the combination and the MIC of compound A (or B) when tested alone, according to Equation 2. Assays were performed in 3 biological replicates.

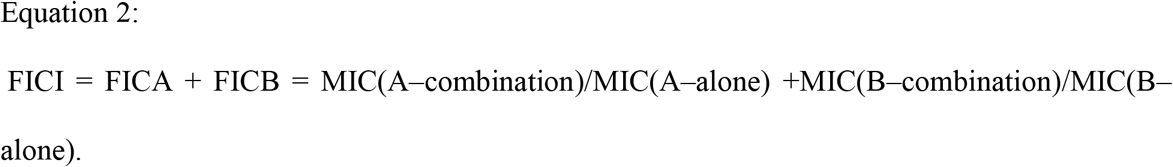

## Acknowledgments

T.P.S.C. would like to thank the National Health and Medical Research Council of Australia (APP1091976) and Australian Research Council (DE190100806) for fellowship and funding support. M.A.P. acknowledges the Australian Research Council for funding support (DP150103313). This research was supported by the Defence Science Institute, an initiative of the State Government of Victoria, with a scholarship awarded to J.A.W., who is also the recipient of a Research Training Program scholarship. C.K.G. is a recipient of the La Trobe University Postgraduate Research Scholarship and La Trobe University Full Fee Research Scholarship. We would also like to acknowledge the La Trobe University Comprehensive Proteomics Platform for providing infrastructure and expertise.

## Author Contributions

T.P.S.C., J.M.S. and M.A.P. designed experiments; T.P.S.C., J.A.W. and C.K.G. performed experiments and analysed data; M.D.H. and B.M.A. provided reagents and infrastructure; T.P.S.C. wrote the manuscript; all authors provided revisions and edits.

